# Endogenous tau released from human *ReNCell VM* cultures by neuronal activity is phosphorylated at multiple sites

**DOI:** 10.1101/2024.06.02.597022

**Authors:** Ghadir Sindi, Sazan Ismael, Reaz Uddin, Kira G. Slepchenko, Robert A. Colvin, Daewoo Lee

## Abstract

Tau is an intracellular protein but also known to be released into the extracellular fluid. Tau release mechanisms have drawn intense attention as these are known to play a key role in Alzheimer’s disease (AD) pathology. However, tau can also be released under physiological conditions although its physiological function and release mechanisms have been poorly characterized, especially in human neuronal cells.

We investigated endogenous tau release in *ReNCell* VM, a human neuroprogenitor cell line, under physiological conditions and found that tau is spontaneously released from cells. To study activity-dependent release of endogenous tau, human *ReNCell VM* culture was stimulated by 100μM AMPA or 50mM KCl for one-hour, tau was actively released to the culture medium. The released tau was highly phosphorylated at nine phosphorylation sites (pSites) detected by phospho-specific tau antibodies including AT270 (T175/T181), AT8 (S202/T205), AT100 (T212/S214), AT180 (T231), and PHF-1 (S396/S404), showing that these pSites are important for activity-dependent tau release from human *ReNCell* VM. Intracellular tau showed various phosphorylation status across these sites, with AT270 and PHF-1 highly phosphorylated while AT8 and AT180 were minimally phosphorylated, suggesting that AT8 and AT180 pSites exhibit a propensity for secretion rather than being retained intracellularly. This activity-dependent tau release was significantly decreased by inhibition of GSK-3β, demonstrating that GSK3β-dependent phosphorylation of tau plays an important role in its release by neuronal activity.

In this study, we showed that *ReNCell* VM serves as a valuable model for studying endogenous physiological tau release. Further, *ReNCell* model can be also used to study pathological release of human tau that will contribute to our understanding of the progression of AD and related dementias.

**Highlights:**

- Activity-dependent release of endogenous human tau from human ReNCell VM cultures occurs under physiological conditions.
- Released human tau is phosphorylated at nine sites (pSites) in the proline-rich domain and the C-terminal domain detected by AT270 (T175/T181), AT8 (S202/T205), AT100 (T212/S214), AT180 (T231), and PHF-1 (S396/S404) tau antibodies, strongly suggesting that these pSites are important for activity-dependent tau release from human ReNCell VM.
- In contrast, intracellular human tau proteins have different phosphorylation status among these nine pSites: AT270 and PHF-1 pSites are highly phosphorylated, but AT8 and AT180 are weakly phosphorylated, suggesting AT8 and AT180 pSites are release-sensitive phosphorylation motifs.
- Activity-dependent release of endogenous human tau is decreased by a tau kinase GSK-3β inhibitor SB 216763, indicating that GSK-3β-dependent phosphorylation plays an important role in activity-dependent tau release.
- The human *ReNCell* culture is an excellent model system to study mechanisms underlying physiological release of endogenous tau.

## Introduction

Tau protein is encoded by microtubule-associated protein tau (MAPT) gene. The function of tau is to bind microtubules and maintain their stability in axons. In line with tau’s function, it is localized through cytosol. However, several studies demonstrated that tau can be released under both physiological and pathological conditions (Yamada et al, 2011; Pooler et al, 2013; Merezhko et al, 2020). The released tau is known to play a key role in the propagation of tauopathies (Wu et al, 2016; Merezhko et al, 2020), but the physiological function of extracellular (or released) tau has been poorly characterized.

The most common post-translational modification of tau is phosphorylation (Neddens et al., 2018; Wegmann et al, 2021; Xia et al, 2021). Tau phosphorylation at specific pSites influences its normal function, leading to microtubule destabilization (Guo et al., 2017). Additionally, neurofibrillary tangle (NFT) is a hallmark of Alzheimer’s Disease (AD) and other tauopathies. NFT are composed of hyperphosphorylated tau aggregates (Noble et al, 2013; Simic et al, 2016). Many tau pSites identified in healthy individuals were found hyperphosphorylated in AD patients (Barthèlemy et al., 2019; Wegmann et al., 2021), suggesting phosphorylation plays a crucial role in AD. Phosphorylation of tau is also known to be a key contributor for its release (Merezhko et al, 2018; Perez et al, 2018). The main kinase that phosphorylates tau is glycogen synthase kinase-3β (GSK-3β), which phosphorylates tau at 42 sites (pSites), and 29 of these GSK-3β specific sites are observed in AD brains (Martin et al., 2013; Guo et al, 2017). However, further exploration is required to understand the role of phosphorylation in both physiological and pathological tau release from neurons.

Interestingly, studies have shown that tau can be released by neuronal activity under physiological conditions (Pooler et al, 2013; Yamada et al, 2014) although the functional role of activity-dependent release of tau is not known. It has been shown that neuronal excitability increases during the early stages of AD (Busche et al, 2008; Siskova et al, 2014; Hall et al, 2015). For example, mild cognitive impairment in AD correlates with hyperactivity in the hippocampus (Dickerson et al, 2005; Busche et al, 2008). Therefore, the impact of activity-dependent release of phosphorylated tau is of significant interest to understand both physiological and pathological roles of released tau.

In this study, we used a cell culture model to study activity-dependent tau release using immortalized human neuronal cell line *ReNCell VM* (Donato et al., 2007), which can be differentiated into neurons and glial cells. We observed that endogenous dimeric tau is spontaneously released from human *ReNCell VM* cells. Under neuronal stimulation with AMPA and KCl, tau dimer release was significantly increased in addition to activity-dependent monomeric tau release. We also described the phosphorylation profiles of intracellular and released tau at nine disease-associated pSites known to be phosphorylated by GSK-3β (Amaral et al., 2021) using five phospho-specific antibodies: T175/T181 (AT270), S202/T205 (AT8), T212/S214 (AT100), T231 (AT180), and S396/S404 (PHF-1). Our study shows that tau phosphorylation plays a critical role in tau release and these nine pSites are similarly important for activity-dependent tau release from human *ReNCell VM*.

## Results

### *ReNCell VM* human neural progenitor cells (hNPCs) express endogenous tau

To study activity-dependent tau release, a human neural progenitor cell *ReNCell VM* was used (Donato et al, 2007). First, *ReNCell VM* was differentiated for 14 days using the standard method of differentiation by the removal of growth factors from culture media (Donato et al, 2007). Then, tau (+) cells in differentiated *ReNCell* culture were stained with anti-human tau (hTau) antibody (HT7; Fig. 1A). A significant number (37.84%) of undifferentiated *ReNCells* (first day in culture) were positive for tau, and this number increased over time (Fig. 1C). We found that the number of tau (+) cells reached a plateau at 9 days of differentiation (Fig. 1C). At 14 days of differentiation, 80.70% ±7.40 of cells were tau (+) (Fig. 1B). In this study, ReNCell cultures differentiated for 14 days were used to examine activity-dependent human tau (hTau) release, from now on these cells will be called differentiated ReNcell culture.

**Figure 1.**
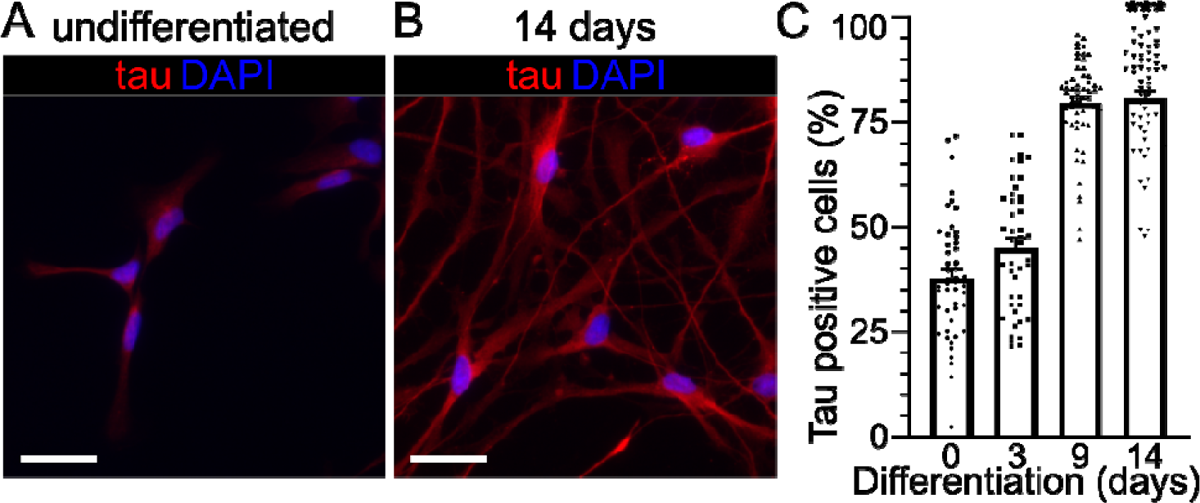
Human tau (hTau) expression in human *ReNCell* VM neuronal culture. **(A and B)** Two images showing cultured cells that were stained with anti-human total Tau (DAKO K9JA) antibody (red) at 0 (undifferentiated) and 14 days in vitro (DIV). The scale bar represents 50μm. **(C)** Tau (+) signals were quantified relative to DAPI signals (blue) in the same field of view at 0 (undifferentiated), 3, 9 and 14 DIV (n=3). Error bars indicate mean ± SEM. Student t-test, ***p <0.001.

### Neuronal depolarization induces tau release from *ReNCell VM* culture

Using differentiated *ReNCell VM* cultures, we examined whether endogenous human tau (hTau) can be released by neuronal depolarization. Differentiated *ReNCell VM* cultures were exposed for one hour to a glutamate receptor agonist AMPA and high KCl, both of which induce neuronal depolarization. Conditioned media (CM) was collected, and tau was concentrated by immunoprecipitation and quantified with ELISA. We found that both 100μM AMPA and 50mM KCl significantly increased hTau release compared to unstimulated control (Fig. 2A). This shows that hTau release from *ReNCell VM* is increased by neuronal activity. We confirmed that the enhancement in hTau release was not associated with changes in hTau level inside the cells because intracellular hTau levels were similar between control and AMPA/KCl groups (Fig. 2B). To determine the integrity of cells after neuronal stimulation, lactate dehydrogenase (LDH) assay was performed. No significant difference was observed in LDH activity in cells stimulated with 50mM KCl compared with control cells (Supplementary Fig. S1A). These results demonstrate that hTau is released as a result of neuronal activity and not from damaged cells. Since 50mM KCl increased hTau release about two-fold greater than 100mM AMPA (Fig. 2A), we used 50mM KCl exposure for the rest of the experiments.

**Figure 2.**
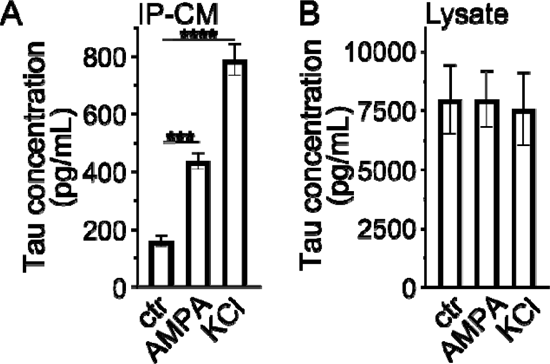
Depolarization-dependent hTau release from differentiated human *ReNCell*. **(A)** ELISA analysis showing released hTau in conditioned media (CM). ReNCell cultures were treated with 100uM AMPA and 50mM KCl for 1 hr. Tau release was normalized to total protein in cell lysate. Data from 3 independent experiments. **(B)** Intracellular hTau levels were not altered by AMPA and KCl stimulation. Error bars indicate mean ± SEM. Student t-test, *p <0.05.

### Biochemical characterization of hTau in *ReNcell VM* culture

To further characterize the biochemical properties of released hTau protein, cell lysate and CM were harvested after 50mM KCl or physiological KCl (5.6 mM – control) exposure from differentiated ReNcells. To concentrate released tau, CM was immunopurified (IP-CM) and resolved on western blots. We found that one-hour exposure to 50mM KCl significantly increased the intensity of hTau bands compared with the control (Fig. 3A & B), which confirmed ELISA results (Fig. 2A).

**Figure 3.**
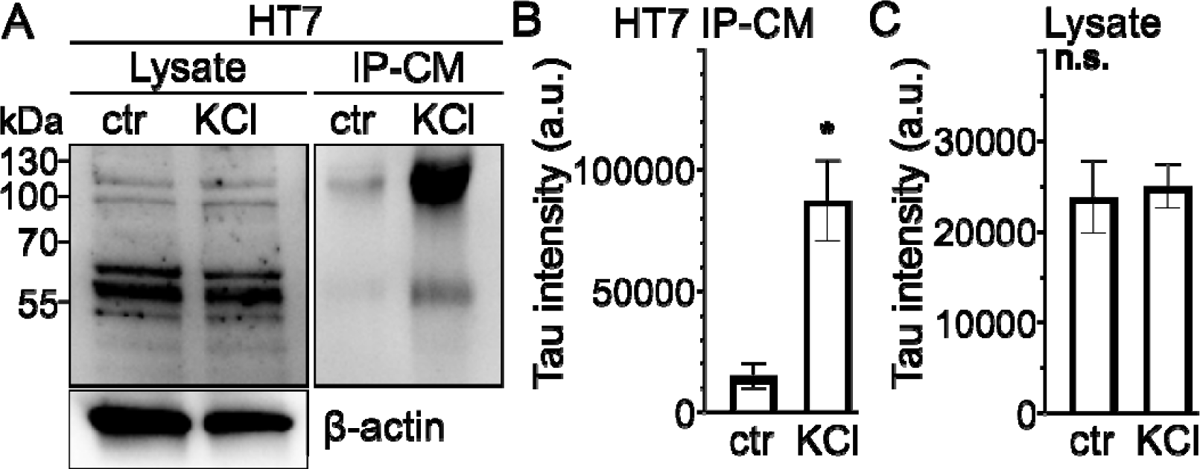
Intracellular and released hTau from human *ReNCell* culture. **(A)** A representative western blot (WB) image showing intracellular tau in lysate and released tau immunopurified from conditioned media (IP-CM). Tau was released to the culture media after 50mM KCl stimulation for 1 hr at 14 DIV. (**B and C**) Tau bands quantified in CM and lysate were between 50 and 120kD. Quantification of HT7 intensity in IP-CM from control and KCl treated sample groups (B). (C) shows quantification of intracellular hTau in cell lysate. Data from 3 independent experiments. Student t-test, *p<0.05.

Several hTau bands were observed in the range of <50 to >120 kDa inside the cell. Lower molecular weight bands most likely correspond to monomeric and/or truncated hTau, while higher molecular weight bands are expected to be dimerized hTau (Fig. 3A). Two bands that were in the range of 50 and 120 kDa were detected in CM suggesting that activity-dependent released tau is monomeric and dimeric (Fig. 3A & B). In addition, an increase in hTau release was not associated with changes in hTau level inside the cell because the intensity of tau bands in lysates from control and 50mM KCl treated cells were similar (Fig. 3C)

### Phosphorylation profile of intracellular hTau and its release from *ReNcell VM*

Several studies have shown that tau phosphorylation is essential for microtubule stabilization and tau release (Saman et al., 2012; Guo et al., 2017; Ismael et al., 2021). We next examined the phosphorylation profiles of intracellular and released hTau using five well-characterized phospho-specific antibodies (Fig. S3): AT270 (T175/T181), AT8 (S202/T205), AT100 (T212/S214), AT180 (T231) which recognize pSites in the proline-rich domain, and PHF-1 (S396/404) which recognizes pSites in the C-terminal domain. We used differentiated *ReNcell* cultures after physiological KCl (control) or 50mM KCl exposure for one hour. Cell lysate and CM were harvested for western blot analysis. AT270, AT100, and PHF-1 antibodies detected multiple bands inside the cell in the range of ∼50 - 60 and ≥130 kDa (Fig. 4A-E). In contrast, anti-AT8 signals were barely observed (Fig. 4B) and anti-AT180 was not detected at all in cell lysate (Fig. 4C). It was noticeable that AT100 selectively detected strong tau bands of molecular weight higher than 100 kD while PHF-1 detected mainly lower molecular weight bands (<70 kD). These results indicate that intracellular hTau was phosphorylated at different pSites depending on its structure (i.e., monomer versus dimer/oligomer). In addition, intracellular hTau is weakly phosphorylated at AT8 (S202/T205) and AT180 pSites (T231). In contrast, regardless of the phospho-specific antibodies used, hTau proteins released by neuronal stimulation were detected at 55 and 100 kD bands. They were phosphorylated at AT270, AT8, AT100, AT180 and PHF-1 epitope pSites (Fig 4A-E). Next, we investigated unphosphorylated pSites using TAU-1 antibody which is known to recognize pSites of S195, S198, S199, and/or S202 in the proline-rich domain (Tatebayashi et al, 2002). We observed that intracellular hTau at lower molecular weight bands (around 55 kDa) was unphosphorylated (Fig. 5 A). Interestingly, released hTau was found unphosphorylated in both 55 and 100 kDa bands (Fig. 5A). The intensity of hTau-1 bands in IP-CM was significantly increased after 50mM KCl treatment (Fig. 5A & B). There was no difference in intracellular unphosphorylated tau (Fig. 5C).

**Figure 4.**
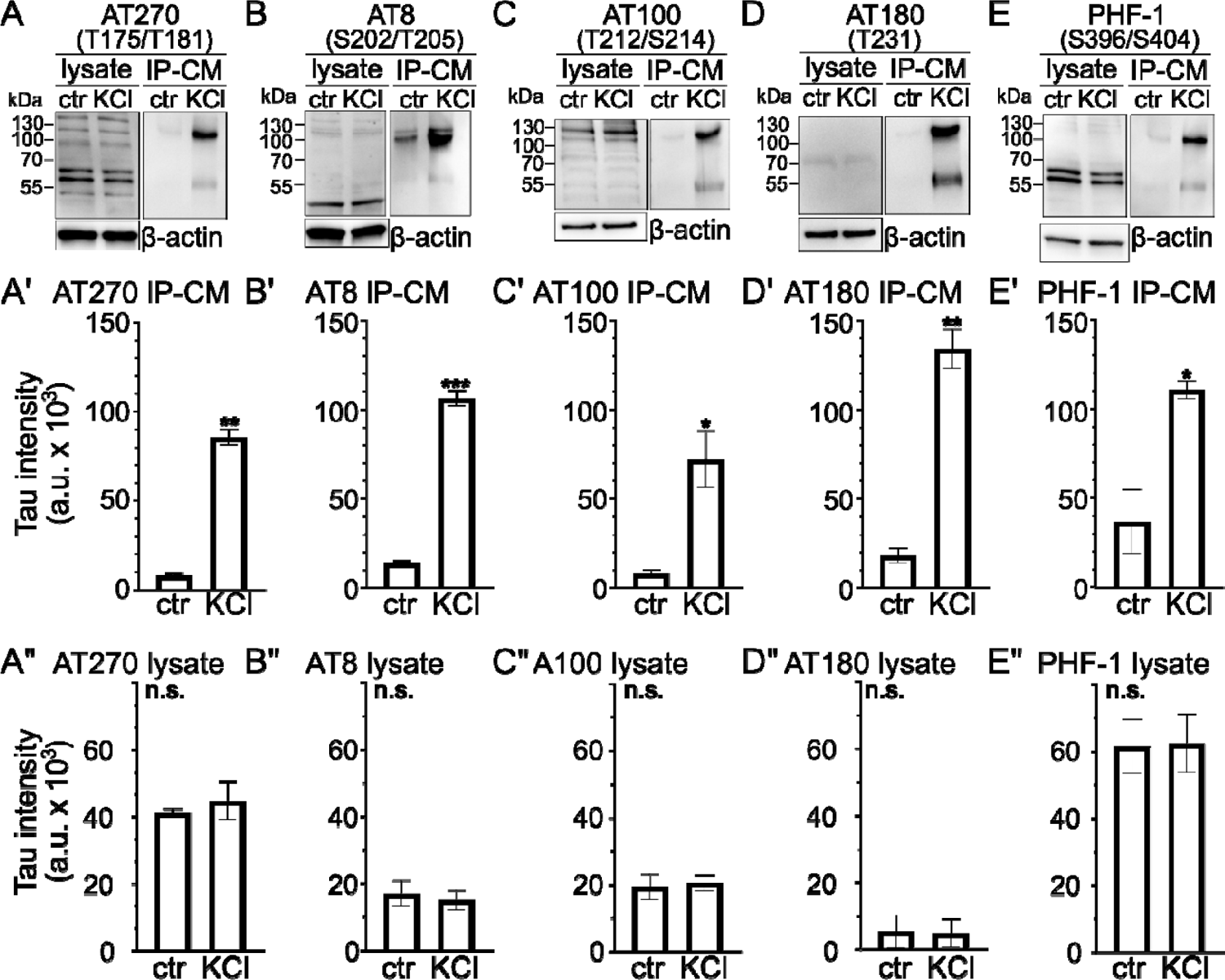
Phosphorylation profiles of intracellular and released hTau from *ReNCell*. **(A-E)** Representative western blot images showing phosphorylated hTau from cell lysate and IP-CM at pSites recognized by AT270 (A), AT8 (B), AT100 (C), AT180 (D), and PHF-1 (E). **(A’-E’)** Quantification of phosphorylated hTau released to CM at pSites detected by AT270 (A’), AT8 (B’), AT100 (C’), AT180 (D’), and PHF-1 (E’). **(A”-E”)** Quantification of phosphorylated hTau from cell lysate at pSites detected by AT270 (A”), AT8 (B”), AT100 (C”), AT180 (D”), and PHF-1 (E”). Data from 3 independent experiments. Tau bands between 50 and 100kDa were quantified relative to β-actin in cell lysate. Error bars indicate Mean ± SEM. Student t-test, *p<0.05, **p<0.01, ***p<0.001.

**Figure 5.**
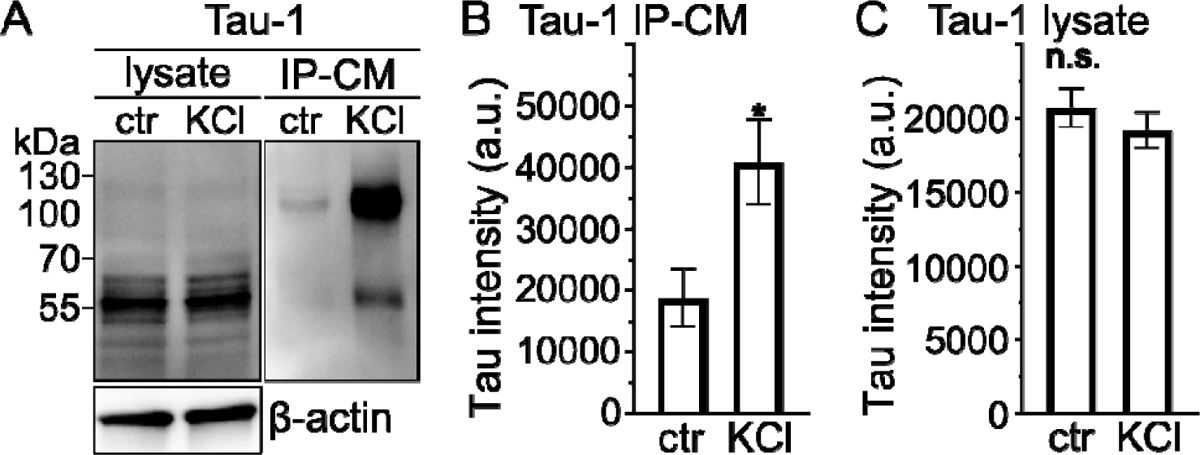
Dephosphorylation of release hTau from *ReNCell* culture at TAU-1 sites. **(A)** A representative western blot (WB) image probed with TAU-1 antibody showing intracellular tau in lysate and released tau immunopurified from conditioned media (IP-CM). Tau was released to the culture media after 50mM KCl stimulation for 1 hr at 14 DIV. (**B and C**) Tau bands quantified in lysate and CM were between 50 and 120kD. Quantification of HT7 intensity in IP-CM from control and KCl treated sample groups (B). (C) shows quantification of intracellular hTau in cell lysate. Data from 3 independent experiments. Student t-test, *p<0.05.

Our results show that released hTau is phosphorylated at multiple pSites but some pSites are apparently not phosphorylated (Figs. 4 & 5). To determine whether phosphorylated or unphosphorylated tau is selectively released by neuronal activity, we calculated fold changes of released tau (Table 1). Tau release was increased more than 7-fold by neuronal activity when it was detected by total tau antibody HT7 (not phospho-specific). All pSites signals in PRD (i.e., AT270, AT8, AT100, AT180) were also similarly increased while PHF-1 signal in C-terminal was decreased compared to total tau HT7 signal. In addition, the fold change of TAU-1 unphosphorylated signal was lower than total hTau (HT7) and phosphorylated hTau released (Table 1). There was no difference in phosphorylation status of intracellular hTau detected by AT270, AT8, AT100, AT180 and PHF-1 with and without neuronal stimulation (Fig. 4). Overall, released hTau is phosphorylated at multiple pSites but also not phosphorylated at TAU-1 pSites. However, phosphorylation does not increase activity-dependent hTau release under physiological conditions as fold increase detected by total tau antibody HT-7 and phospho-specific antibodies was similar (Table 1). In contrast, fold changes detected by TAU-1 and PHF-1 were significantly reduced, indicating that dephosphorylation of hTau at TAU-1 and PHF-1 pSites decreased subsequent release (Table 1). Our results showed that tau phosphorylation plays an important role in activity-dependent hTau release from *ReNCell* cultures.

**Table 1.** Fold changes of Western blot hTau signals from cell lysate and IP-CM by neuronal stimulation. Fold changes were calculated as the ratio of hTau bands measured after 50mM KCl versus control (5.6mM KCl) for each antibody.

| ***Tau epitope*** | Cell lysate | IP-CM |
| --- | --- | --- |
| HT7 | 1.08 | 7.13 |
| AT270 | 1.08 | 10.72 |
| AT8 | 0.91 | 7.75 |
| AT100 | 1.10 | 9.12 |
| AT180 | 1.12 | 8.13 |
| PHF-1 | 1.01 | 4.36* |
| TAU-1 | 0.93 | 2.27** |

### GSK-3β activity regulates activity-dependent hTau release from *ReNCell VM*

GSK-3β is a major tau kinase that phosphorylates tau. However, the role that GSK-3β activity plays in activity-dependent hTau release is unknown. In this study, we investigated whether GSK-3β inhibition influenced activity-dependent hTau release. We used a potent, selective GSK-3β inhibitor SB216763 (SB; Wagman et al, 2004). This inhibitor demonstrated a protective mechanism against cell death in primary neuronal culture (Cross et al., 2001). In addition, it has been reported that SB showed promising therapeutic activity in decreasing AD symptoms (Ruiz & Eldar-Finkelman, 2022).

Differentiated *ReNCell* cultures were randomly divided into two groups: cultures were treated with 10μM SB (for 2 hours) or DMSO (control). Then, both culture groups were treated with 50mM KCl (for one hour) to induce neuronal depolarization. After KCl treatment, cell lysate and CM were harvested for western blot analysis. We showed that intracellular total hTau (HT7) significantly decreased after 10μM SB compared with control cells (Fig. 6A & C). In addition, GSK-3β inhibition reduced the release of total hTau compared with control cells (Fig. 6A & B). To determine if SB treatment causes cell damage, lactate dehydrogenase (LDH) assay was used. No significant difference was observed in LDH activity after SB treatment compared to control (Fig. S1B). Since intracellular hTau level was decreased after SB treatment, another total hTau antibody was used to confirm this reduction by SB treatment. We stained the membrane with another total tau K9JA (DAKO) antibody (Wang et al, 2009). K9JA epitope is in microtubule repeats and the C-terminal domain (aa 243-441), and thus shows no preference for phosphorylated or unphosphorylated hTau. hTau bands were observed in the range of ∼50 to 120 kDa in cell lysate (Fig. 6 D). hTau positive bands in CM revealed that monomers and dimers of hTau were released via neuronal stimulation (Fig. 6D & E). Released hTau bands from IP-CM were reduced after SB treatment compared with untreated cells (Fig. 6E). Consistent with HT7 antibody, K9JA antibody showed a decrease in total intracellular hTau after SB treatment (Fig. 6D &F).

**Figure 6.**
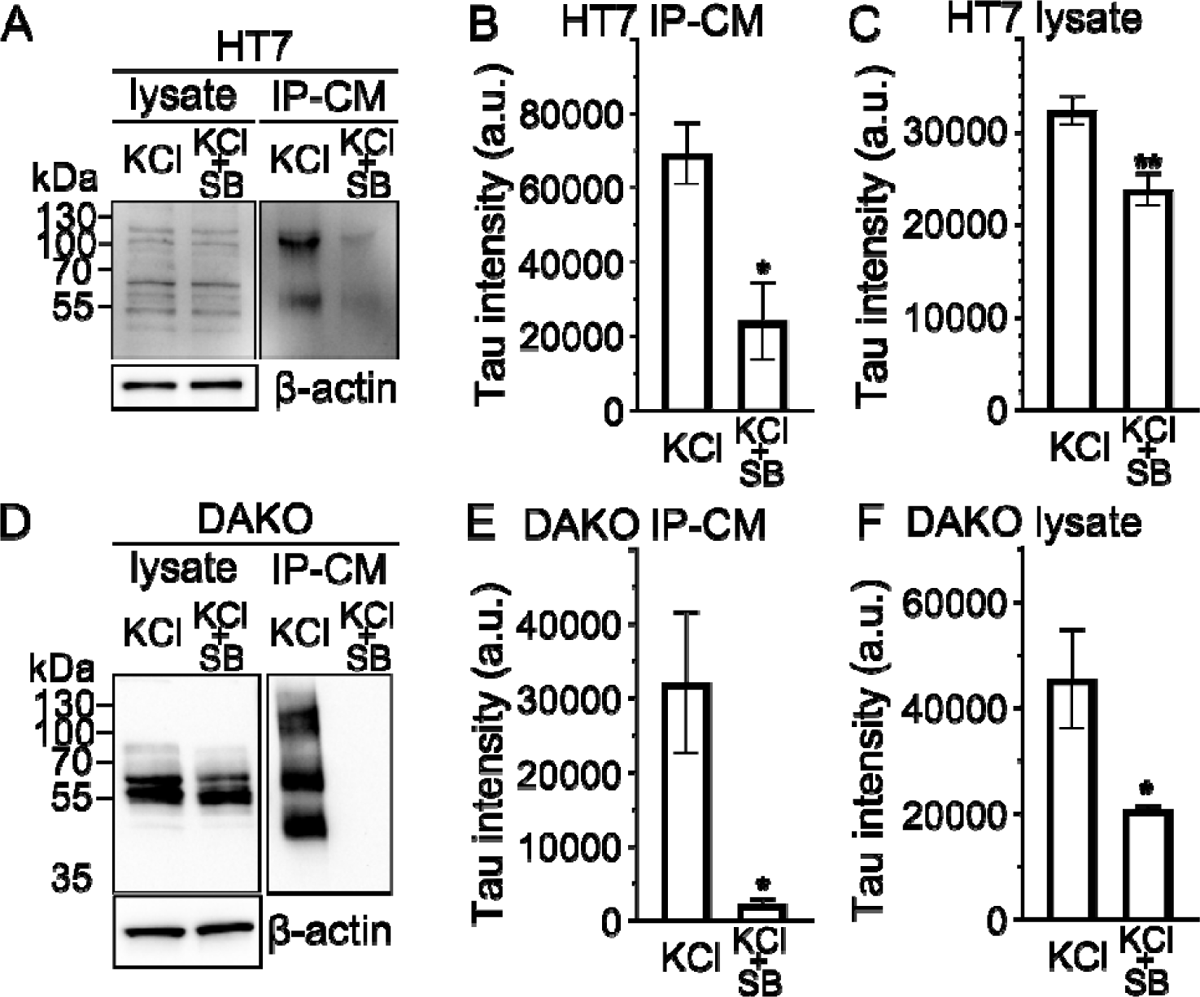
GSK-3β inhibition influences activity-dependent hTau release from ReNCell. **(A)** Representative western blot image showing that total hTau (probed by HT7) from cell lysate and IP-CM is decreased after SB216763 (SB) treatment (10μM for 2hrs). **(B and C)** Quantification of hTau bands in cell lysate and IP-CM, respectively. HT7 antibody was used to probe hTau bands. **(D)** Western blot image showing total hTau (probed by K9JA) from cell lysate and IP-CM decreases after SB treatment (10μM for 2hrs). **(E and F)** Quantification of hTau bands in cell lysate and IP-CM, respectively. hTau bands between 50 and 120kDa were quantified relative to β-actin in cell lysate. Data from 3 independent experiments. Error bars indicate Mean ± SEM. Student t-test, *p <0.05, **p<0.01.

### Phosphorylation levels of released hTau at specific pSites are decreased after GSK-3β inhibition

In this experiment, we studied the role of nine individual pSites in activity-dependent hTau release by inhibiting GSK-3β with 10μM SB for 2 hours. Differentiated *ReNCell* cultures were treated with either SB or DMSO. As in previous experiments, we stimulated cells with 50mM KCl treatment, and collected lysate and CM for analysis on western blots. Results showed that phosphorylation levels of intracellular hTau at pSites of T175/T181 (AT270), T212/S214 (AT100), S396/S404 (PHF-1) were significantly reduced after SB treatment (Fig 7A, C, E).

**Figure 7.**
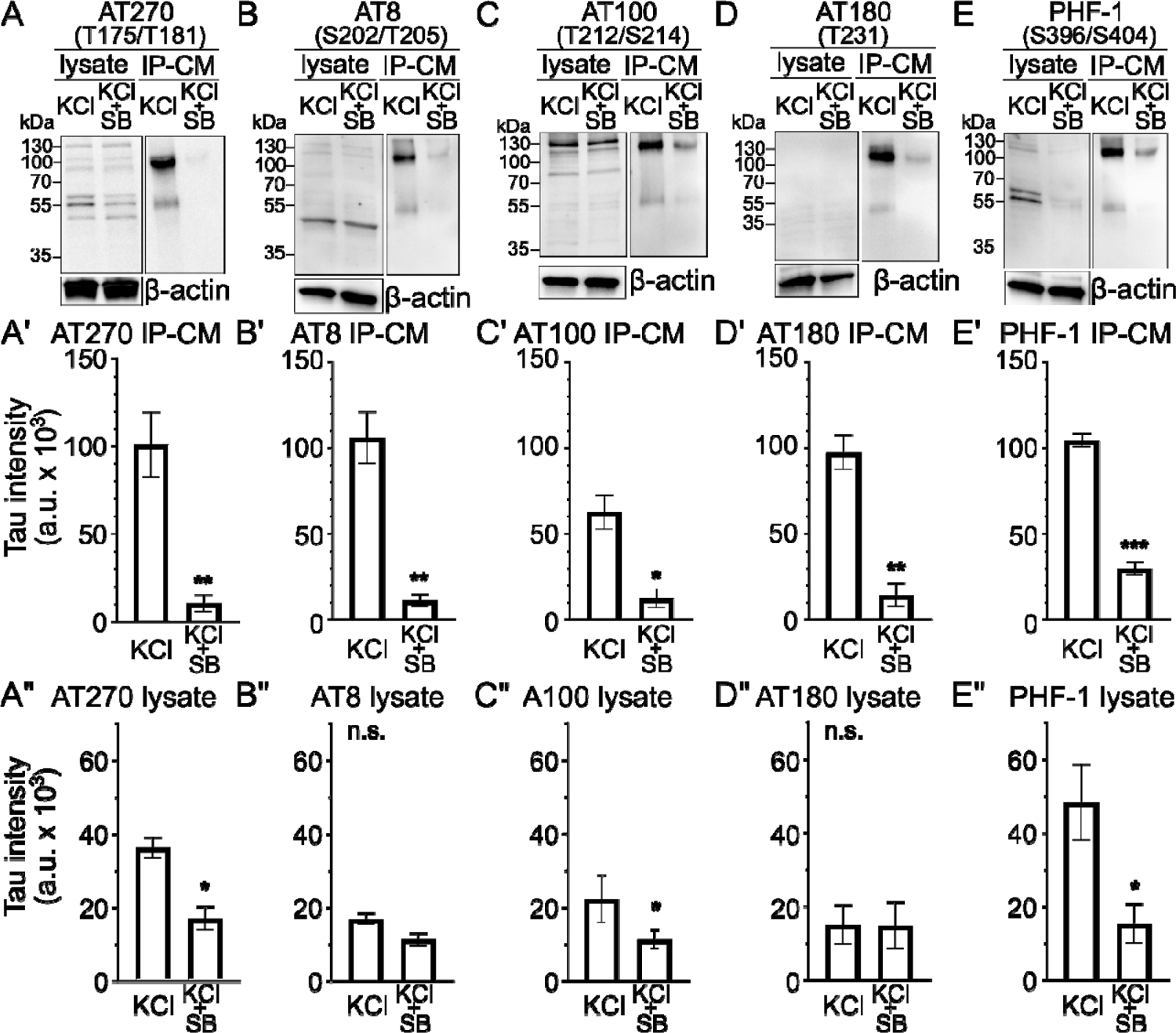
Phosphorylation levels of released hTau at specific pSites are decreased after GSK-3β inhibition. *ReNCell* cultures (14 DIV) were incubated with SB (10μM) for 2hrs before 50mM KCl stimulation (1 hour). **(A-E)** Representative western blot images showing phosphorylated hTau from cell lysate and IP-CM at pSites recognized by AT270 (A), AT8 (B), AT100 (C), AT180 (D), and PHF-1 (E). **(A’-E’)** Quantification of phosphorylated hTau released to CM at pSites detected by AT270 (A’), AT8 (B’), AT100 (C’), AT180 (D’), and PHF-1 (E’). **(A”-E”)** Quantification of phosphorylated hTau in cell lysate at pSites detected by AT270 (A”), AT8 (B”), AT100 (C”), AT180 (D”), and PHF-1 (E”). Data from 3 independent experiments. Tau bands between 50 and 120kDa were quantified relative to β-actin in cell lysate. Error bars indicate Mean ± SEM. Student t-test, *p<0.05, **p<0.01, ***p<0.001.

However, phosphorylated hTau at pSites of S202/T205 (AT8) and T231 (AT180) was not changed (Fig. 7B and D). This is mainly because AT8 and AT180 pSites are not phosphorylated in intracellular tau proteins in control groups.

We also examined phosphorylation status of released tau in the presence of GSK3β inhibitor (SB). The intensity of tau bands in IP-CM detected by AT270, AT8, AT100, AT180, and PHF-1 were significantly decreased after SB treatment compared with control (Fig. 7A-E). Interestingly, the fold decrease of phosphorylated hTau bands in IP-CM was not different from that for total hTau HT-7 antibody (Table 2). In summary, our results show that tau phosphorylation at multiple pSites by GSK3β is important for activity-dependent tau release under physiological conditions.

**Table 2.** Fold changes (log_2_) of Western blot hTau signals from cell lysate and IP-CM by neuronal stimulation after SB treatment. Fold changes were calculated as the ratio of hTau bands measured after SB treatment versus control for each antibody.

| ***Tau epitope*** | Cell lysate | IP-CM |
| --- | --- | --- |
| HT7 | -0.44 | -1.94 |
| AT270 | -1.16 | -3.79 |
| AT8 | -0.60 | -3.32 |
| AT100 | -0.95 | -2.82 |
| AT180 | -0.01 | -3.32 |
| PHF-1 | -1.78** | -1.82 |

## Discussion

In this study, we explored the physiological release of endogenous tau using the human neuroprogenitor cell line *ReNCell* VM. During neuronal stimulation, endogenous tau was actively released into the culture medium. The released tau was phosphorylated at multiple phosphorylation sites (pSites). Additionally, GSK3β inhibition decreased both activity-dependent tau release and its phosphorylation levels, underscoring the importance of GSK3β-dependent tau phosphorylation in influencing tau release. Our results show that the human *ReNCell* culture is an excellent model system to study activity-dependent physiological release of endogenous human tau and the mechanisms underlying the role of phosphorylation in its release.

Under physiological conditions, the release of tau has been established. Microanalysis techniques and ELISA assays demonstrated the detection of released wild-type tau in both the interstitial fluid (ISF) and cerebrospinal fluid (CSF) in the mouse brain (Yamada 2017; Xia et al., 2021). Additionally, investigations in mouse primary neuronal cultures, as documented by Karch et al. (2012), further support the extracellular detection of tau. Previous studies, such as those conducted by Pooler et al. (2013) and Yamada et al. (2014), have underscored that endogenous tau release is a normative physiological process in the brain. Notably, neuronal activity has been identified as an inducer of tau release under physiological conditions, occurring independently of cell death (Pooler et al., 2013; Yamada et al., 2014; Yamada, 2017). These findings strongly imply a physiological role for extracellular tau, although the precise physiological functions of extracellular tau remain largely unknown. In addition, several studies showed that extracellular tau binds and activates the M1 and M3 muscarinic acetylcholine receptors (Gomez-Ramos et al., 2008; Pernègre et al., 2019). Finally, Morozova et al (2019) showed that uptake of normal tau in neurons promotes the formation of neuronal processes whereas a pseudo-phosphorylated variant known as pathological human tau (PH-Tau) disrupts them.

The physiological function of tau is based on different post-translational modifications, mainly phosphorylation (Mietelska-Porowska et al., 2014; Guo et al., 2017). Throughout this study, we examined the phosphorylation profile of intracellular tau at nine pSites in the proline-rich domain and C-terminal region: T175/ T181 (AT270), S202/T205 (AT8), T212/S214 (AT100), T231 (AT180), and S396/S404 (PHF-1). We found that intracellular tau was phosphorylated at AT270, AT100, and PHF-1 pSites. However, pSites recognized by AT8 and AT180 epitopes were weakly phosphorylated inside the cells. Consistent with our results, it has been demonstrated that intracellular tau was phosphorylated at AT270 (T181), AT100, and PHF-1 in Hela cells (Plouffe et al., 2012). In addition, our recent findings showed that AT8 and AT180 pSites were less phosphorylated inside the cell in *Drosophila* primary neuronal culture (Ismael et al., 2021). However, Plouffe et al. (2012) claimed that intracellular tau at AT8 and AT180 sites was phosphorylated. These differences might be due to the different cell types and/or tau isoforms used in each study, but it prompts us to further investigate phosphorylation profiles of intracellular tau under physiological conditions.

Our results showed that released tau proteins were phosphorylated at AT270, AT8, AT100, AT180, and PHF-1 pSites. We demonstrated that tau release was enhanced by neuronal depolarization with KCl and AMPA. This agrees with our recent findings that the release of phosphorylated hTau at AT8, AT100, AT180, and PHF-1 pSites was increased from *Drosophila* neurons by neuronal activity (Ismael et al., 2021). Furthermore, released tau from M1C cells, derived from human neuroblastoma, overexpressing wild-type tau was phosphorylated at AT270, AT8, AT100, and AT180 (Saman et al., 2012). Karch et al. (2013) also showed that phosphorylated tau at pSites T181 and S396 was released from human neuroblastoma cells (SH-SY5Y). Consistent with our findings, a recent study demonstrated that released tau from human iPSCs was phosphorylated at T181 (AT270), T231(AT180), and S396 (PHF-1) (Wadhwani et al, 2019). It is interesting to note that some of these pSites are also phosphorylated in tau in the cerebrospinal fluid (CSF) of AD brains (Hampel et al, 2005; SuárezLJCalvet et al, 2020; Janelidze et al., 2020; Ashton et al, 2022). Tau phosphorylated at T181, T217, or T231 rises in CSF in the initial stages of preclinical AD (SuárezLJCalvet et al., 2020; Ashton et al, 2022), indicating that these pSites can be useful biomarkers for the early detection of AD. In addition, Barthélemy et al (2023) showed that CSF tau phosphorylated at T205 and T217 can be reliable biomarkers of amyloid and tau pathology in AD as these two pSites are well correlated with tau PET measures. All these strongly support the importance of revealing the role of specific pSites to understand the physiological and pathological roles of released tau.

Glycogen synthase kinase-3 beta (GSK-3β) is a crucial kinase responsible for phosphorylating various target proteins, including tau. In our investigation, we employed - 216763 (SB) to inhibit GSK-3β, a compound known for its potent inhibitory effects on GSK-3β activity (Coghlan et al., 2000). Previous studies have illustrated SB’s ability to attenuate tau phosphorylation in the postnatal rat brain (Selenica et al., 2007). Our study demonstrated that treatment with SB significantly decreased activity-dependent tau release from *ReNCells,* along with a reduction in intracellular tau levels within the cells compared to the control cells that were not treated with SB. Specifically, SB treatment led to a significant decrease in intracellular tau phosphorylated at T175/T181 (AT270), T212/S214 (AT100), and S396/S404 (PHF-1).

Moreover, tau signals phosphorylated at nine pSites were decreased in SB-treated cells compared to controls, highlighting the importance of these sites in activity-dependent tau release. These findings are supported by Cross et al. (2001), who observed reduced tau phosphorylation at T181 (AT270) and S202 (AT8) following SB treatment in cerebellar granule neurons. Furthermore, SB treatment inhibited extracellular α-synuclein-induced tau hyperphosphorylation at S396 (Gąssowska et al., 2014) and reduced tau phosphorylation at S396/S404 in clonal CHO cells expressing human tau (Boom et al., 2009). However, conflicting results have been reported regarding SB’s effect on tau hyperphosphorylation at S202 in human neuroblastoma SH-SY5Y cells (Wisessaowapak et al., 2021). Nevertheless, our study underscores the significant role of GSK-3β-mediated tau phosphorylation in physiological tau release.

Our data showed that intracellular and released tau from *ReNCells* was also dephosphorylated at TAU-1 sites (S195, S198, S199, and/or S202). To determine the contribution of phosphorylation and dephosphorylation of tau on its release, we quantified the fold increase of phosphorylated versus dephosphorylated tau by neuronal stimulation (Table 1). The fold increase of released total tau was probed by the phospho-independent tau antibody (HT7). Our results showed that there was no significant difference in the fold increase between total tau HT7 and all phospho-specific antibodies (AT270, AT8, AT100, and AT180) except PHF-1. However, the fold increase of released tau phosphorylated at these pSites was significantly higher than dephosphorylated tau recognized by TAU-1 (Table 1). Our results support that released tau is phosphorylated at multiple pSites. However, phosphorylated tau was not preferentially released under physiological conditions. In contrast, several studies showed that phosphorylated tau is preferentially released in a pathological condition (Plouffe et al, 2012; Katsinelos et al, 2018; Ismael et al, 2021). Overall, our findings demonstrate the significance of tau phosphorylation in its activity-dependent release. While our results indicate that released tau is phosphorylated at multiple pSites, our data from Table 1 suggest that under physiological conditions, phosphorylation is not required for increased tau release. This stands in stark contrast to its role under pathological conditions. In a previous study, we demonstrated a preferential release of phosphorylated tau (Ismael et al., 2021). Tau phosphorylation appears to play a crucial role in its release under both physiological and pathological conditions. However, it is noteworthy that phosphorylated tau is selectively released in pathological scenarios, likely facilitating neurodegeneration in Alzheimer’s disease (AD). This underscores the significance of hyperphosphorylation as a pivotal factor in the cell-to-cell propagation of tau pathology.

While extracellular tau plays a crucial physiological role, its pathological implications have been extensively examined. Disruption of neuronal activity is involved in AD pathogenesis. Studies have shown that neuronal excitability increases during the early stages of AD (Busche et al, 2008; Siskova et al, 2014; Hall et al, 2015). For example, mild cognitive impairment in AD correlates with hyperactivity in the hippocampus (Dickerson et al, 2005; Busche et al, 2008).

Therefore, the impact of activity-dependent release of pathological tau on tauopathy is of significant interest. There is considerable focus on the cell-to-cell propagation of tau facilitated by neuronal activity. Through advanced optogenetic and chemogenetic interventions, Wu et al. (2016) showed that elevating neuronal activity not only amplifies the release and transfer of tau in laboratory settings but also intensifies tau pathology in living organisms. We also showed that human tau release is increased by optogenetic-induced depolarization from *Drosophila* primary neuronal culture (Ismael et al., 2021). Despite intensive studies, the molecular underpinnings of release and propagation of pathological tau proteins remain to be explored. Tau phosphorylation also plays a pivotal role in pathological functions of tau (Spillantini & Goedert, 2013; Tenreiro et al, 2014; Simic et al, 2016). Particularly, phosphorylation is crucial for tau propagation, including its release (Plouffe et al, 2012; Merenzhko et al, 2018). In our investigation, we observed phosphorylation across all nine hTau pSites examined, consistent with findings from a study conducted in Hela cells, where mimicking phosphorylation at 12 sites associated with AD led to an increased tau release (Plouffe et al, 2012). Additionally, recent research by Katsinelos et al (2018) demonstrated that abnormally phosphorylated tau is released more efficiently than dephosphorylated tau. Similarly, Wadhwani et al. (2019) identified phosphorylated tau at pSites T181, T231, and S396 in endogenous tau released from neurons derived from human induced pluripotent stem cells (PSCs), with phosphorylation at S199 notably absent, aligning with previous literature (Pooler et al, 2013; Mohamed et al, 2014; Croft et al, 2017), supporting the notion that tau released spontaneously or by neuronal activity is phosphorylated at multiple pSites. However, contrary evidence suggests that released tau may be dephosphorylated or hypo-phosphorylated. For instance, Pooler et al. (2013) detected dephosphorylated endogenous tau using western blot analysis with antibodies specific to dephosphorylated tau (TAU-1) or phosphorylated tau (PHF-1). Mohamed et al. (2014) also observed dephosphorylation in tau released from resting mouse neurons. Notably, however, these studies employed a limited number of phospho- or dephospho-specific antibodies (e.g., PHF-1, Tau-1), resulting in an incomplete phosphorylation profile of released tau.

How does tau detected in *ReNcells* compare to tau found in human brain? Human cortex lysates, probed with HT7 antibody (Ercan et al. 2017; Shultz et al. 2015; Zhou et al. 2018) and DAKO antibody (Ercan et al. 2017) showed similar tau patterns in *ReNcells*, probed with the same antibodies. The phosphorylation status of tau in human brain and *ReNcells* is also comparable, because cortical homogenates from healthy and Alzheimer’s human brains show analogous tau phosphorylation as we detected in *ReNcells*, using antibodies AT8, AT270, AT100 and PHF1 (Zhou et al. 2018). Interestingly the phosphorylation of tau in *ReNcells* is closer to patterns of phosphorylation seen in human brains with Alzheimer’s (Zhou et al. 2018). Therefore, there are comparable tau isoforms and phosphorylation in the human brain and *ReNcells*, suggesting that these cells are a suitable cellular model to study tau and tau release.

In summary, the *ReNCell* VM model emerges as a valuable tool for investigating activity-dependent tau release under both physiological and pathological conditions. Our findings emphasize the significant role of GSK-3β inhibition in modulating activity-dependent tau release, as evidenced by the reduction in tau release following such inhibition. Additionally, we observed that phosphorylation sites (pSites) play a crucial role in regulating activity-dependent tau release, indicating that tau release is influenced both by neuronal activity and phosphorylation. Investigating activity-dependent tau release provides valuable insights into strategies for mitigating the progression of tauopathies, including AD. Our study also suggests the potential for designing novel therapies targeting GSK-3β and pSites to suppress tau release and in turn diminish its propagation between neurons. Consequently, the human *ReNCell* culture serves as a model system for elucidating the mechanisms underlying the physiological release of endogenous human tau. In addition, because we showed spontaneous and activity-dependent endogenous tau release from the *ReNCell* culture, we suggest that these cells can be a good model for pathological tau release, if the cells are stressed with conditions known to cause AD, like oxidative stress, neuroinflammation and others. Known risk factor genes can be expressed in ReNcells and using these cells as a model will contribute to our understanding of the progression of AD and related dementia as has been done using *ReNCell* expressing human mutant APP and PSEN1 genes (Choi et al, 2014; Park et al, 2018).

### Experimental procedures

#### ReNCell VM Culture

*ReNCell VM* (Millipore-sigma, cat# SCC008) culture protocol was prepared as described previously (Donato et al, 2007; Cristóvaõ et al, 2012). Laminin (Fisher Scientific, cat# CC0955MG) was diluted in DMEM/F12 and used to coat the tissue culture flasks. Cells were plated in “complete medium” that was made of DMEM/F12 (Hyclone, cat# SH30004.02), neural cell supplement B27 (Fisher Scientific, cat# 17-504-044), 10 U/ml heparin sodium salt (Sigma Aldrich, cat# H3149), GlutaMAX (Fisher Scientific, cat# 35-050-061), 50μg/ml gentamicin (Fisher Scientific, cat#1 5-750-060), 10ng/ml basic fibroblast growth factor (bFGF) (PeproTech, cat# AF-100-188), and epidermal growth factor (EGF) (PeproTech, cat# AF-100-15). Cell cultures were maintained in the incubator until they became 80-90% confluent. Subculture was done with Accutase enzyme (Fisher Scientific, cat# MT25058CI). Cells were differentiated by the standard method (stdD) (Donato et al, 2007; Cristóvaõ et al, 2012). This differentiation method depends on the removal of growth factors from the culture medium. On the first day, cells were plated on laminin-coated coverslips and allowed to grow in the complete medium. On the next day, growth factors were removed from the medium and replaced with the maintenance medium (complete medium without growth factors) to start differentiation. The maintenance medium was changed every day for the first 6 days and then every other day to keep differentiated cells for 14 days in vitro (DIV).

#### AMPA and KCl Treatment

100μM AMPA (Tocris, cat#0254), 50mM KCl and 5.6mM physiological KCl were prepared. Differentiated *ReNCells* (14DIV) were treated for one hour at 37°C with either 5.6mM KCl (control), 100μM AMPA, or 50mM KCl to induce neuronal depolarization. To quantify secreted tau, conditioned media (CM) was collected, centrifuged at 10,000 rpm for 10 minutes, and then used for immunoprecipitation assay (IP) (see details in “Immunoprecipitation Assay”). For quantification of intracellular tau, cells were harvested for cell lysate preparation (see details in “Lysate preparation”).

#### Kinase Inhibitor Treatment

A selective GSK-3β inhibitor SB 216763 (Tocris, cat# 1616) was prepared at 1mM concentration (stock solution) and diluted in *ReNCell VM* culture medium at a final concentration of 10μM. SB stock was dissolved in DMSO and stored at −20°C. Differentiated *ReNCell VM* culture was treated with SB for two hours at 37°C.

#### Immunofluorescence Assay (IFA)

An immunofluorescence assay was performed as previously described (Ismael et al, 2021). For tau staining, anti-human total Tau (Agilent DAKO, cat# 2004; 1:2000) was incubated overnight at 4°C. Then, coverslips were incubated with a secondary antibody for one hour (FITC or tetramethylrhodamine labeled; Invitrogen). Lastly, coverslips were mounted on a slide glass and neurons were examined under the fluorescence microscope (Olympus 1X71). Spot CCD digital camera was used for taking images (Darya et al., 2009). Images were taken from 5-10 random areas per one coverslip. Positive signals and DAPI signals were counted as previously described (Wiemerslage et al, 2013).

#### Lysate Preparation

Radio-immunoprecipitation (RIPA) assay buffer (Thermo Fisher Scientific, cat# 89900) was used to homogenize cells. Phosphatase/protease inhibitor cocktail (Thermo Fisher Scientific, cat# 78440) was added to the lysis buffer. The lysate was mixed on a shaker for one hour at 4°C and then centrifuged at 10,000 rpm for 10 minutes. The supernatant was collected and stored at −80°C and used for ELISA and Western blot experiments. To normalize tau in western blot lysate samples, β-actin band intensity was used.

#### Immunopurification (IP)

IP was performed using Pierce crosslink immunoprecipitation kit (Thermo Scientific, cat# 26147). Antibodies of interest (HT7, Fisher Scientific, cat# MN1000; K9JA, Agilent DAKO, cat#A0024) were crosslinked to protein A/G plus agarose. The column was made by adding protein A/G agarose slurry inside the column. Then, the column was washed two times with 1x coupling buffer (CB) at 3,500 rpm for 1 minute. The slurry was then incubated with (10μg) of antibody at room temperature for one-hour. Then, it was washed with CB three times at 3,500 rpm for 1 minute. After that, the beads were incubated with (50μl) Disuccinimidyl suberate (DSS) (2.5μl 20X CB, 38.5μl distilled water, and 2.5mM DSS diluted in DMSO) at room temperature for one hour. Next, the beads were washed with elution buffer (pH 2.8) three times, and with IP lysis buffer twice at 3,500 rpm for 1 minute. Finally, the column was stored in 1x CB at 4°C to be used later for IP.

#### IP was started by incubating CM with pierce control agarose resin at 4°C for one hour

Then, CM was centrifuged and washed with IP lysis buffer and incubated in the column that had antibody-crosslinked beads (HT7 or DAKO) overnight at 4°C. Next day, the column was centrifuged and washed with IP lysis buffer two times and with condition buffer one time at 3,500 rpm for 1 minute. After that, the immunopurified protein was eluted with 60μl elusion buffer and the column was centrifuged at 3,500 rpm for one minute and the flow through was collected and stored at −80°C to be used for ELISA and Western blot experiments.

#### Enzyme-Linked Immunosorbent Assay (ELISA)

Human Tau ELISA kit (Abcam, cat# ab229394) was used according to manufacturer protocol. A fluorescence microplate reader (BioTek Synergy HTX platereader) was used for reading the signals (Excitation 530nm/ Emission 590nm). Data were analyzed using GraphPad prism and normalized to lysate protein in the same cell culture.

#### Western Blotting

Lysate and IP-CM were loaded equally into 12% sodium dodecyl sulfate-polyacrylamide gel (SDS-PAGE; Bio-Rad cat# 4561043). Next, a nitrocellulose membrane (Pall, cat# 66489) was used to transfer proteins. The membrane was blocked with 3% bovine serum albumin (3% BSA) that was diluted in tris-buffered saline plus tween 20 (TBS-T) for 2 hours at room temperature. Then, the membrane was incubated with primary antibody overnight at 4°C.The next day, the membrane was washed with TBS-T three times for 10 minutes and then incubated with secondary antibody for 2 hours at room temperature. The membrane was washed three times for 15 minutes. Protein bands were detected using chemiluminescent substrate.

#### Statistical Analysis

Data are shown as mean ± SEM. Data are represented by at least three independent repeats of each experiment. Each repeat was a biological replicate. Means were compared using an unpaired two-tailed Student’s *t*-test. p-values are * < 0.05, ** < 0.01, *** < 0.001.

## Acknowledgements

This work was partially supported by NIH R15AG065925 and R21NS130256. We would like to thank Aeran Lee for helping with *ReNCell* neuronal cultures and western blot experiments.

## Author contribution

GS and DL designed the experiments. GS performed most experiments and analyzed the results. RU performed a part of western blot experiments. KS made final figures. KS and RAC provided valuable inputs to earlier versions of manuscripts and final edits. RAC also analyzed LDH assay results. GS and DL wrote the final version of the manuscript.

