## Supplementary figures for "Endogenous tau released from human *ReNCell VM* cultures by neuronal activity is phosphorylated at multiple sites"

**Figure S1. Lactate dehydrogenase assay.** Absorbance of control, KCl, KCl+SB and max (maximum release). Error bars indicate Mean ± SEM. Each dot represents a biological replicate.

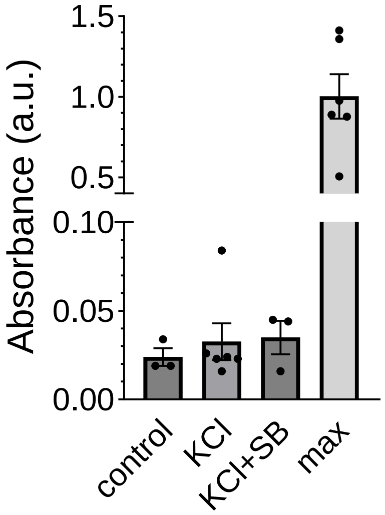

**Figure S2. The western blot of ReNcell lysates and conditioned media (CM) that was immunoprecipitated (IP-CM).** The whole blot is presented in this figure.

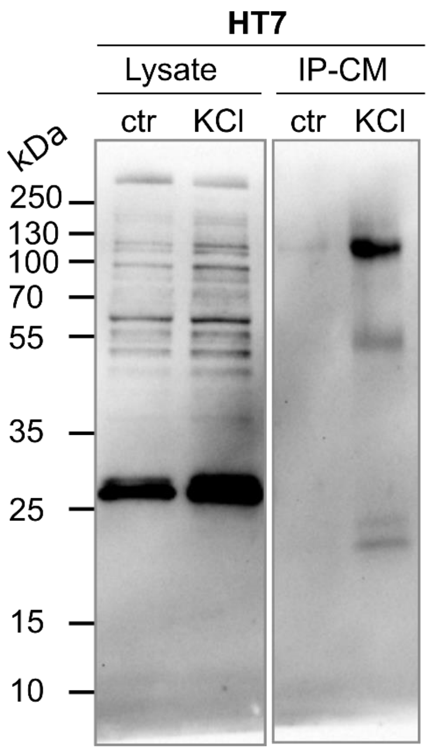

**Figure S3**. **A diagram of antibodies and epitopes used in this study.** The antibody and epitopes are listed in the full-length human tau with different regions labeled. Arrows represent each antibodies' epitope.

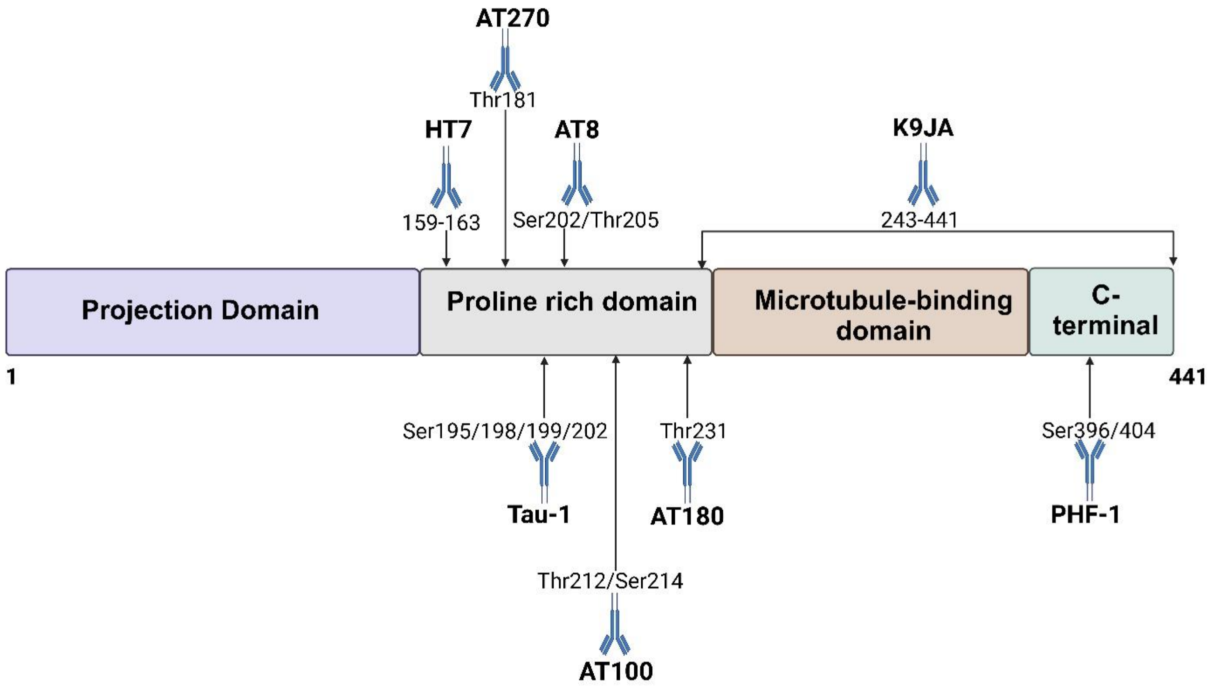
